# Losing Control: Sleep Deprivation Impairs the Suppression of Unwanted Thoughts

**DOI:** 10.1101/813121

**Authors:** Marcus O. Harrington, Jennifer E. Ashton, Subbulakshmi Sankarasubramanian, Michael C. Anderson, Scott A. Cairney

## Abstract

Unwanted memories often enter conscious awareness when we confront reminders. People vary widely in their talents at suppressing such memory intrusions; however, the factors that govern suppression ability are poorly understood. We tested the hypothesis that successful memory control requires sleep. Following overnight sleep or total sleep deprivation, participants attempted to suppress intrusions of emotionally negative and neutral scenes when confronted with reminders. The sleep-deprived group experienced significantly more intrusions (unsuccessful suppressions) than the sleep group. Deficient control over intrusive thoughts had consequences: whereas in rested participants suppression reduced behavioural and psychophysiological indices of negative affect for aversive memories, it had no such salutary effect for sleep-deprived participants. Our findings raise the possibility that sleep deprivation disrupts prefrontal control over medial temporal lobe structures that support memory and emotion. These data point to an important role of sleep disturbance in maintaining and exacerbating psychiatric conditions characterised by persistent, unwanted thoughts.

## Introduction

Memories of unpleasant experiences and thoughts can intrude into conscious awareness when we confront reminders to them. Individuals suffering from psychiatric conditions such as post-traumatic stress disorder (PTSD) and major depressive disorder (MDD) typically experience a disproportionate number of unwanted memory intrusions, and difficulties in limiting the duration and recurrence of these intrusions compound negative mood and affective dysregulation (Brewin, Gregory, Lipton, & Burgess, 2010; Mihailova & Jobson, 2018; Moritz et al., 2014; Newby & Moulds, 2011; Payne, Kralj, Young, & Meiser-Stedman, 2019). Our capacity for inhibiting intrusive thoughts might therefore play a fundamental role in maintaining mental health and wellbeing (Gagnepain, Hulbert, & Anderson, 2017).

The ability to control intrusive memories and thoughts can be studied in the laboratory by measuring people’s success at suppressing memory retrieval when confronted with reminders to unwanted thoughts. For example, one widely used task, known as the Think/No-Think (TNT) paradigm (Anderson & Green, 2001), requires participants to either actively engage (‘Think’) or suppress (‘No-Think’) memory retrieval when presented with reminder cues to associated memories, often aversive images, An intrusion occurs when participants’ attempts to suppress retrieval during ‘No-Think’ trials fail, and the reminder cue triggers an involuntary retrieval of the associated memory. Reminder cues typically elicit an intrusion less than 50 percent of the time (Gagnepain et al., 2017; Hellerstedt, Johansson, & Anderson, 2016; Levy & Anderson, 2012; van Schie & Anderson, 2017), but people vary widely in memory control ability (Levy & Anderson, 2008). The factors contributing to this variability are poorly understood. Identifying determinants of successful retrieval suppression could contribute to our understanding of vulnerability to disorders characterised by intrusive thoughts (Brewin et al., 2010; Moritz et al., 2014; Streb, Mecklinger, Anderson, Johanna, & Michael, 2016).

Retrieval suppression ability is thought to be intrinsically linked to inhibitory control. According to the inhibitory deficit hypothesis (Levy & Anderson, 2008), individual differences in regulating intrusive memories originate from variation in underlying inhibition function. This hypothesis predicts that conditions that strain inhibitory control will likewise undermine the ability to suppress unwanted thoughts. In healthy adults, sleep deprivation impairs cognitive functioning (Alhola & Polo-Kantola, 2007; Walker, 2009; Wild, Nichols, Battista, Stojanoski, & Owen, 2018), especially executive control (Drummond, Paulus, & Tapert, 2006; Nilsson et al., 2005), making sleep loss an important candidate factor mediating fluctuations in thought control. Indeed, mental fatigue arising from sustaining an effortful task can increase the frequency of intrusions during the TNT task (van Schie & Anderson, 2017). Moreover, the prefrontal – medial temporal lobe (MTL) networks involved in retrieval suppression (Benoit, Hulbert, Huddleston, & Anderson, 2015; Gagnepain et al., 2017; Levy & Anderson, 2012) are disrupted by sleep loss (Yoo, Gujar, Hu, Jolesz, & Walker, 2007), suggesting that losing sleep may heighten people’s vulnerability to intrusive thoughts (Chee, 2004; Mazur, Pace-Schott, & Hobson, 2002; Thomas et al., 2000, 2003; van Schie & Anderson, 2017; Yoo et al., 2007). Chronic sleep disturbance is also a formal symptom of most psychiatric conditions, particularly PTSD (Maher, Rego, & Asnis, 2006) and MDD (Riemann, Berger, & Voderholzer, 2001).

These observations led us to consider whether the established link between psychiatric conditions and disturbed sleep may be mediated partially by sleep deficits compromising a person’s ability to regulate emotion by suppressing retrieval of aversive thoughts. For example, suppressing retrieval of aversive scenes reduces people’s emotional response to those scenes later on, as revealed by changes in subjective affect ratings for the suppressed stimuli, and the relationship of those changes to prefrontally-driven downregulation of the amygdala during memory intrusions (Gagnepain et al., 2017). This impact of retrieval suppression on perceived emotion (hereinafter referred to as affect suppression) suggests that retrieval suppression contributes to affective homeostasis by reducing the negative tone of unpleasant events. If sleep loss compromises memory control, it may diminish the impact that retrieval suppression has on affective responses to unwanted thoughts, a possibility consistent with the well-documented negative consequences of sleep loss on mood (Dinges et al., 1997; Short & Louca, 2015; Zohar, Tzischinsky, Epstein, & Lavie, 2005). Whether sleep loss disrupts affect suppression arising from memory control, and whether affect suppression effects are mirrored in psychophysiological reactivity to suppressed thoughts, has never been examined.

To determine whether sleep deficits could contribute to the pathogenesis and maintenance of intrusive symptomatology, we investigated the impact of sleep deprivation on memory control in healthy adults. Participants learned associations between faces and emotionally negative or neutral scenes before an overnight interval of sleep or total sleep deprivation (Figure 1a). The following morning, participants completed a TNT task for the face-scene associations (Figure 1b). On each trial of this task, a face was presented alone in either a green or red frame, instructing participants to either actively retrieve (‘Think’) or suppress (‘No-Think’) the associated scene, respectively. Attempts to suppress retrieval of the scene often initially fail, leading the scene to intrude into participants’ awareness involuntarily, despite efforts to stop it. Because involuntary retrievals in the TNT paradigm are (as in real life) unobservable events private to the individual, it was necessary to identify their occurrence through intrusion reports. After each trial, participants reported whether the associated scene had entered awareness on a three-point scale (“never”, “briefly”, “often”). We then quantified variations in memory control success by using the proportion of ‘No-Think’ trials that triggered any awareness of the associated scene, reflecting a momentary failure of retrieval suppression (i.e. reports of “briefly” or “often”; hereinafter referred to as intrusions).

**Fig. 1.**
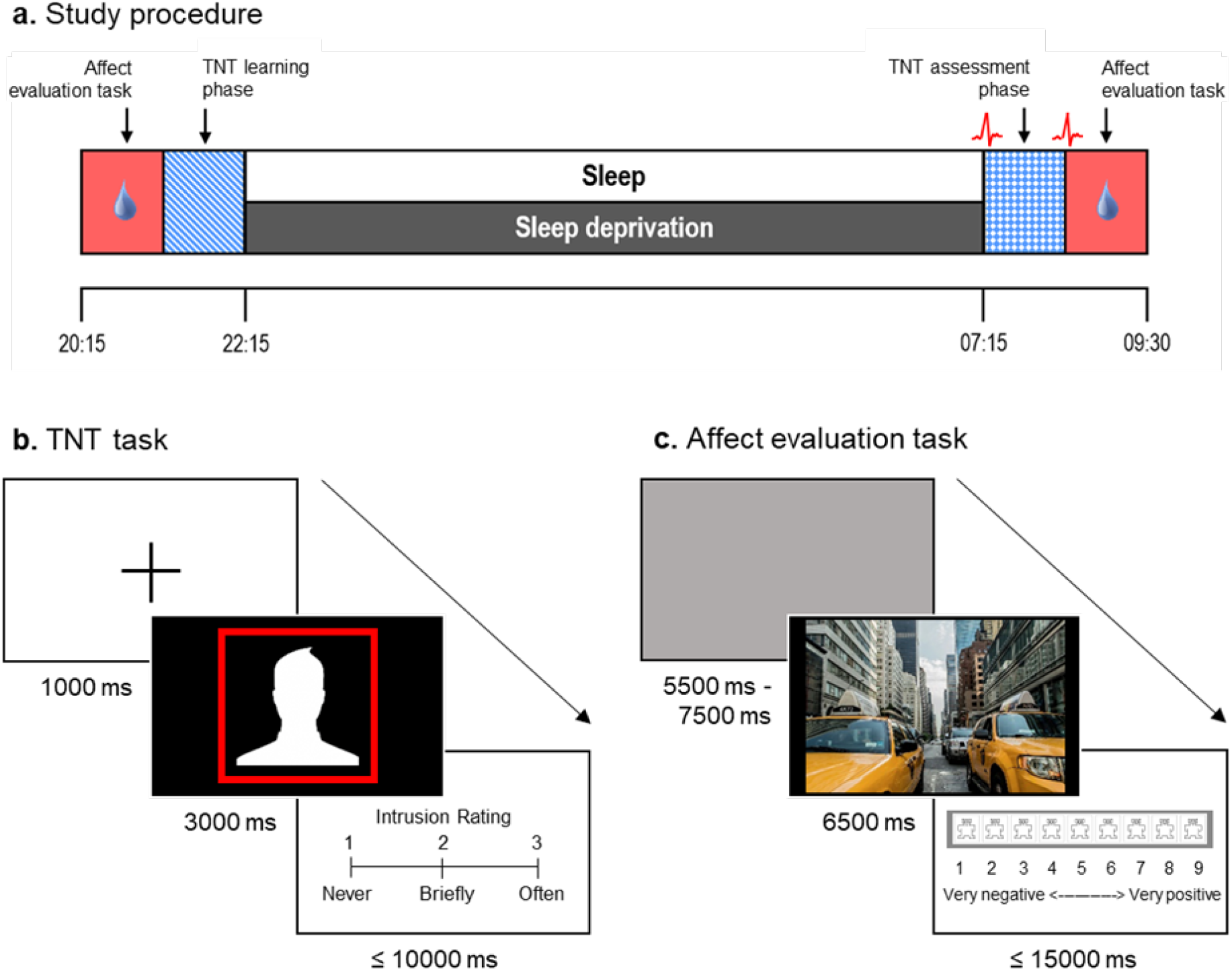
Study procedures and tasks. (a) All participants completed session one in the evening, which included the first of two affect evaluation tasks and the learning phase of the Think/No-Think (TNT) task. Participants in the sleep group slept in the sleep laboratory for ~8 hours. Participants in the sleep deprivation group remained awake across the night. All participants then completed session two, which included the assessment phase of the TNT task and the second affect evaluation task. We collected skin conductance responses (SCRs) to scene presentations throughout the affect evaluation tasks. We recorded resting heart rate variability (HRV) before and after the TNT assessment phase. (b) In the critical TNT assessment phase, we presented participants with faces in red or green frames. For red framed faces (‘No-Think’ trials), we instructed participants to avoid thinking about the associated scene without generating diversionary thoughts; for green framed faces (‘Think’ trials), we instructed them to visualise the associated scene. After each trial, participants reported the extent to which they thought about the paired scene (“never”, “briefly” or “often”). We considered reports of “briefly” or “often” as intrusions. Note that the human face stimuli example has been replaced for the purpose of this preprint. (c) In the affect evaluation task, participants viewed negative and neutral scenes and we asked them to provide an emotional rating for each image on a scale from 1 (very negative) to 9 (very positive).

Intrusion reports provide a validated index of involuntary retrieval. Using intrusion reports, fMRI studies have established that in rested individuals intrusion trials (unsuccessful suppression attempts) trigger greater activation in the right dorsolateral prefrontal cortex (rDLPFC), greater negative coupling of rDLPFC with both the hippocampus and amygdala, and greater downregulation of activity in the latter structures, consistent with a reactive engagement of top-down inhibitory control to suppress the intrusive content (Benoit et al., 2015; Gagnepain et al., 2017; Levy & Anderson, 2012). EEG studies, moreover, establish that reports of intrusions are associated with a brief increase in ERP markers of working memory storage (Hellerstedt et al., 2016), followed by their rapid elimination, consistent with a reactive purging of the intrusion. In contrast, trials which attract intrusion reports of “never” (successful suppression attempts) instead show significant increases in beta frequency power, consistent with successful proactive control of retrieval (Castiglione, Wagner, Anderson, & Aron, 2019). Using these intrusion indices, we predicted that sleep-deprived participants would report more intrusions for ‘No-Think’ scenes than would participants who slept and exhibit impaired ability to reduce the frequency of intrusions over time.

Studies that employ the TNT paradigm often measure the consequences of thought suppression for memory accessibility via a recall test administered after the TNT assessment phase. These studies have shown that suppressing a memory often impairs its later accessibility (suppression-induced forgetting; for review see: Anderson & Hanslmayr, 2014). However, our objective here was instead to test how sleep deprivation influenced the ability to downregulate unwanted thoughts and, consequently, affect suppression for the intruding content. To investigate how the predicted failures in intrusion control following sleep loss influenced affect suppression, we acquired emotional ratings for the scenes both before the overnight delay and also after the TNT phase the following morning (Figure 1c). We also measured skin conductance responses (SCRs) to scene presentations to examine whether psychophysiological measures of sympathetic arousal mirrored suppression-induced changes in subjective affect. We predicted that retrieval suppression would attenuate both subjective and psychophysiological reactivity to negative scenes in the sleep group. Critically, however, if our hypothesis about the role of sleep loss in the pathogenesis of psychiatric disorders is correct, this salutary effect of suppression on negative affect should be significantly reduced in otherwise healthy participants randomly assigned to our sleep deprivation group.

We also measured the physiological correlates of memory control by collecting heart rate variability (HRV) recordings before and after the TNT assessment phase. Spectral analysis of HRV has previously identified two reliable components: high-frequency HRV (HF-HRV; 0.15-0.40 Hz) and low-frequency HRV (LF-HRV; 0,04-0,15 Hz), Importantly, higher HF-HRV is linked to superior executive functioning (for review, see: Thayer & Lane, 2009), including memory control (Gillie, Vasey, & Thayer, 2014), whereas LF-HRV instead increases with fatigue (Tran, Wijesuriya, Tarvainen, Karjalainen, & Craig, 2009). Given these observations, and given the hypothesized disruption to inhibitory control with sleep deprivation, we predicted that higher HF-HRV would be associated with superior affect suppression and intrusion control in the sleep group, but not the sleep deprivation group. We further predicted significantly higher LF-HRV following sleep deprivation than after a night of sleep, providing physiological confirmation of extreme fatigue.

## Methods

### Participants

Sixty healthy individuals were recruited for this study via an online recruitment system and randomly assigned to a sleep group (*n*=30) or a sleep deprivation group (*n*=30). Of these participants, one was excluded for failing to follow task instructions. The reported data relate to the remaining fifty-nine participants (sleep group: *n*=29, 13 male, mean age = 19.79 years, SD = 1.63 years; sleep deprivation group: *n*=30, 12 male, mean age = 20.20 years, SD = 1.75 years) who participated in return for £40 payment or BSc Psychology course credit (University of York). To reduce demand characteristics, the study was advertised and framed to participants as an investigation into the effect of sleep and sleep deprivation on attention.

Participants reported no history of neurological, psychiatric, attention, or sleep disorders, and they typically rose by 08:00 after at least 6 hours of sleep, as indicated by self-report. Beck depression inventory (BDI-II; Beck, Steer, & Brown, 1996) scores (collected at the end of the experiment) did not differ between groups [sleep group: M = 7.52, SEM = 1.08, sleep deprivation group: M = 5.30, SEM = 0.97; *t*(57) = 1.53, *p* = .13]. We requested that participants refrain from consuming alcohol or caffeine for 24 h prior to the experiment. Wristwatch actigraphy was used to ensure that participants did not nap during the day preceding the overnight phase. Written informed consent was obtained from all participants in line with the requirements of the University of York’s Department of Psychology Research Ethics Committee, who approved the study.

### Stimuli

Forty-eight emotionally neutral face images (half male, half female) served as cues in the TNT task. An equal number of scene images (half negative, half neutral) selected from the International affective picture system (IAPS; Lang, Bradley, & Cuthbert, 2005) served as targets. Negative scenes received lower affect ratings compared to neutral scenes from participants in the current study during the first affect evaluation task [*F*(1,57) = 630.89, *p* < .001, *η_p_*^2^ = .92], demonstrating that the scenes elicited the expected emotional response. Face-scene pairs were created by randomly assigning each face cue to a target scene. Three lists of 16 pairs (8 negative; 8 neutral) were created from the 48 pairs to allow three within-subjects TNT conditions (‘Think’, ‘No-Think’, ‘Baseline’). The assignment of pairs to TNT conditions was counterbalanced across participants for each of the sleep groups. Twelve additional face-scene pairs (6 negative, 6 neutral) were created to serve as fillers that were also used in the practice phases. The same 60 scenes (48 experimental + 12 practice) were used for the affect evaluation tasks, which featured an additional 8 filler scenes (4 negative, 4 neutral).

### Procedure

A schematic representation of the study procedure is shown in Figure 1a. Participants completed two sessions that were separated by an overnight delay containing either sleep or sleep deprivation. At 09:00 on the first day participants collected an actigraphy watch that they wore until 20:15 when they arrived at the University of York’s Emotion Processing and Offline Consolidation (EPOC) laboratory.

#### Affect evaluation task

Participants provided emotion ratings for 68 IAPS scene images using a pictorial scale that ranged from a frowning face on the far-left side of the scale to a smiling face on the far-right side of the scale (Figure 1c). On each trial, a scene was presented for 6.5 s. Participants were asked to focus their attention on the scene for the entire time it was on the screen. Following a 2 s blank screen delay, the affect rating scale was displayed prompting participants to provide their rating on a scale from 1 (corresponding to the frowning face on the far-left side of the scale) to 9 (corresponding to the smiling face on the far-right side of the scale). Participants were told that the extreme left side of the scale should be used for scenes that made them feel *completely* unhappy, annoyed, unsatisfied, melancholic, despaired or bored, and the extreme right side of the scale should be used for scenes that made them feel *completely* happy, pleased, satisfied, contented and hopeful. Participants were given 15 s to provide their affect rating for each picture, although they were asked to respond quickly and spontaneously. The trial terminated once an affect rating had been provided or the 15 s time-limit expired, and participants then viewed a blank screen for 5 s, followed by a fixation cross for a random interval of either 0.5 s, 1 s, 1.5 s, or 2 s, followed by the next scene. This pattern continued until all scenes had been viewed and rated. The affect evaluation task was repeated after the TNT assessment phase. SCRs were recorded throughout the affect evaluation tasks.

#### TNT task

Next, participants completed the learning phase of the TNT task. Participants initially encoded face-scene pairs by studying them for 6 s, one at a time. To reinforce learning and ensure adequate knowledge of the face-scene pairs, participants then completed a test phase. Here, faces were displayed individually for up to 4 s, and participants indicated whether or not they were able to *fully visualize* the associated scene. If the participant indicated that they were able to visualize the associated scene, they were next presented with the correct scene alongside two additional foil scenes that they had seen previously in the learning phase but were not paired with that particular face. Participants were required to select the scene associated with the face. If the participant failed to identify the correct scene, or they indicated that they could not visualize the scene associated with the face, their memory for the face-scene pair was probed again later in the test phase. Regardless of whether or not the participant was able to correctly identify the scene paired with a particular face, the correct face-scene pairing was presented for 3.5 s after each trial. Participants were instructed to use this feedback as an opportunity to reinforce their knowledge of the pairs. The test phase continued until each face-scene pair had been correctly identified once. Following completion of the test phase, a second, identical, test phase was administered to reinforce learning. This overtraining procedure was employed to ensure that participants would experience difficulty in preventing scenes from intruding into consciousness when presented with face cues on ‘No-Think’ trials of the TNT assessment phase.

Before the overnight delay, participants carried out a mock TNT assessment phase. Here, we provided detailed and comprehensive training on the TNT task, allowing participants to practice inhibiting memory intrusions and calibrate their intrusion reports for the critical TNT assessment phase the following morning. In the mock task, participants completed 24 trials (12 ‘Think’, 12 ‘No-Think’) which used 12 filler items (each suppressed or retrieved twice) that were different to those used in the TNT assessment phase proper. Participants then went to sleep or remained awake overnight – details about the overnight delay are provided below.

The following morning, participants completed a memory refresher phase in which the face-scene pairs were presented for 1.5 s per pair, offering participants the opportunity to reinforce their knowledge of the pairs. The TNT assessment phase proper was then administered in five blocks, each lasting approximately 8 min. During each block, two repetitions of 16 ‘Think’ (8 negative and 8 neutral face cues) and 16 ‘No-Think’ (8 negative and 8 neutral face cues) items were presented in pseudorandom order, with the two repetitions of each item appearing at least three trials away from each other. Accordingly, participants completed a total of 320 trials (32 trials x 2 conditions x 5 blocks). Face cues appeared inside a green frame on ‘Think’ trials and inside a red frame on ‘No-Think’ trials. For green-framed faces, participants were instructed to visualize, in as much detail as possible, the scene associated with the face for the entire 3 s that it was on the screen. For red framed faces, participants were instructed to focus their attention on the face for the entire 3 s, but simultaneously prevent the associated scene from coming to mind (Figure 1b). In keeping with earlier work (e.g. Gagnepain et al., 2017). participants were told to accomplish this by making their mind go blank, rather than by replacing the unwanted scene with another image, thought or idea. If the scene came to mind automatically, participants were asked to actively push the scene out of mind.

Immediately after each face cue presentation, participants reported whether the associated scene had entered awareness (during the time that the face was on screen) by pressing a key corresponding to one of three options: “never”. “briefly”, and “often”, Participants were instructed to provide a “briefly” response if the scene had briefly entered conscious awareness at any time during the trial. They were told that “often” responses should be provided if the scene had entered awareness several times during the trial, or once but for a longer period of time than would be considered brief. Participants were given up to 10 s to make this rating; however, they were instructed to provide their rating quickly without dwelling on their decision. Participants moved immediately onto the next trial after providing their intrusion rating, These intrusion ratings were collected to ascertain how competent participants were at suppressing scenes associated with faces for ‘No-Think’ trials. Although we collected intrusion reports on a 3-point scale (“never”, “briefly”, “often”) for every trial, in practice, people rarely give “often” ratings on ‘No-Think’ trials (M = 3.15% of ‘No-Think’ trials in the current study, SEM = 0.62%), For simplicity, we therefore followed prior work by combining “briefly” and “often” responses (Benoit et al., 2015; Gagnepain et al., 2017; Hellerstedt et al., 2016; Levy & Anderson, 2012; van Schie & Anderson, 2017), rendering the judgment binary (intrusion or non-intrusion). Participant responses were followed by a jittered fixation cross lasting between 0.5 and 9 s, before the next face cue was presented.

Note that 16 scenes that were included in the TNT learning phase, and the affect evaluation task, did not appear in the TNT assessment phase. These ‘Baseline’ scenes provided an estimate of generalised changes in affect that could be compared with changes resulting from mnemonic suppression (‘No-Think’ scenes).

#### Recognition task

In the morning, after the second affect evaluation task, memory for all face-picture pairs was examined using a recognition test. The procedure and results for this task are reported in Recognition Task S1.

#### Overnight delay

Electrodes were attached to participants in the sleep group following completion of the first session to allow for overnight polysomnographic recording. Lights were turned out at approximately 22:45 and participants were awoken at 06:45, providing an 8 h sleep opportunity. Before the morning session began, participants were given the opportunity to shower and eat breakfast.

Participants in the sleep deprivation condition remained awake in a university seminar room throughout the overnight delay under the supervision of at least one researcher. Participants were tested in groups of two or three; they were permitted to communicate, read, use the PC, watch TV, or play games. Snacks were provided, but participants were not permitted to consume caffeine. Once the second session began, participants had been awake for > 24 h.

### Equipment and data processing

#### Behavioural

All aspects of the TNT task were written and implemented using Presentation (Neurobehavioral Systems, Albany, CA, USA). The affect evaluation task was administered with E-Prime (Psychology Software Tools, Pittsburgh, PA, USA).

Behavioural tasks ran on a desktop computer and visual aspects were displayed on a flat-screen monitor. Participant responses were collected using the computer keyboard. Behavioural data were analysed using SPSS (IBM Corp., Armonk, NY, USA).

##### Intrusion proportion scores

Intrusion ratings provided during the TNT assessment phase were used to classify each trial as either eliciting an intrusion (“briefly” or “often” responses) or not (“never” responses) in a binary fashion (Benoit et al., 2015; Gagnepain et al., 2017; Hellerstedt et al., 2016; Levy & Anderson, 2012; van Schie & Anderson, 2017). Unwanted retrieval events were separated into “briefly” and “often” to ensure that all intrusions, irrespective of their persistence or strength, were reported. We calculated the proportion of trials that evoked a memory intrusion, separately for each participant, trial block (1-5), TNT condition (‘Think’, ‘No-Think’), and scene valence (neutral, negative). Overall intrusion proportion scores were also calculated by averaging mean intrusion proportion scores across trial blocks, separately for each TNT condition and scene valence.

##### Intrusion slope scores

To measure changes in intrusion frequency for ‘No-Think’ trials across TNT blocks, we calculated proportionalised intrusion slope scores, separately for each participant and scene valence (neutral, negative). Slope scores were calculated by taking the slope of the intrusion frequencies across the five TNT trial blocks. This value was divided by intrusion frequency in the first block, to account for the fact that initial intrusion rates varied and participants with more initial intrusions had more room to decrease their intrusion frequency. We then multiplied the values by −1 to render the (primarily) negative scores positive, with increasingly positive scores reflecting increasing levels of control at downregulating the frequency of intrusions. This measure was z-normalized within the participant’s face-scene pair counterbalancing group (Hellerstedt et al., 2016; Levy & Anderson, 2012). Z-normalizing within each counterbalancing group allows us to quantify a participant’s intrusion slope score with respect to a group of participants receiving precisely the same stimuli in the TNT task.

##### Relapse probability

Each item that appeared in the TNT task was presented 10 times (twice in each TNT block). This repetition allowed us to investigate how well a participant’s efforts to suppress a particular item on one trial carried forward to the next trial involving that same item. The frequency of relapses for ‘No-Think’ items was calculated as the number of ‘No-Think’ trials in repetitions 2-10 where an item elicited an intrusion, but that specific item had been successfully suppressed on its immediately preceding presentation. This was done separately for each participant, trial transition (1 to 2, 2 to 3 … 9 to 10), and scene valence (neutral, negative). Each value was divided by the number of successfully suppressed items (non-intrusions) during the first repetition of that trial transition (e.g. repetition 3 of transition 3 to 4) to produce relapse probability scores (van Schie & Anderson, 2017). This was to account for the fact that individuals with fewer intrusions on the first repetition of each trial transition had greater relapse potential.

##### Affect suppression

Affect ratings gathered during the affect evaluation tasks were used to measure overnight changes in subjective emotional reactivity to the scenes. Mean affect rating values were calculated for each participant, session (pre-delay, post-delay), TNT condition (‘Think’, ‘No-Think’, ‘Baseline’) and scene valence (neutral, negative). Affect suppression scores were then calculated by subtracting the averaged values at session one from those at session two. To account for individual differences in emotional responses at session one, affect suppression scores in each condition were divided by the mean affect rating at session one to produce proportionalised affect suppression scores. Greater scores reflect more positive affect evaluations at session two, as compared to session one.

#### Electrodermal activity

Electrodermal activity recorded during the affect evaluation tasks was used to measure overnight changes in SCRs to the presentation of scenes. In session one and two, the mean SCR was calculated separately for each TNT condition and valence category. Session one SCRs were then subtracted from session two SCRs to generate SCR difference scores (dSCR). Further details about equipment and data processing are available in Methodological Details S1.

#### Heart rate variability

Electrocardiography recordings were administered before and after the TNT assessment phase for HRV analysis. We calculated LF-HRV, which reflects both sympathetic and parasympathetic control over heart rate, and HF-HRV, which reflects parasympathetic or vagal modulation (Laborde, Mosley, & Thayer, 2017).

Further details about equipment and data processing are available in Methodological Details S2.

#### Polysomnography

Polysomnography data was acquired to ensure that participants in the sleep group obtained an adequate amount of sleep, and to allow us to characterise their sleep architecture (see Methodological Details S3 and Table S1).

### Data analysis

Intrusion control measures were analysed for ‘No-Think’ trials with a 2 (Valence: Negative/Neutral) x 2 (Group: Sleep/Sleep Deprivation) mixed ANOVA. Analyses of intrusion proportion and relapse probability included the additional factors Trial Block (1-5) and Trial Transition (1 to 2, 2 to 3 … 9 to 10), respectively. Affect suppression (subjective ratings and SCRs) was analysed for negative scenes with a 2 (TNT Condition: No-Think/Baseline) x 2 (Group: Sleep/Sleep Deprivation) mixed ANOVA. HRV measures were analysed using a 2 (Time: Pre-TNT/Post-TNT) x 2 (Group: Sleep/Sleep Deprivation) ANOVA. Note that ANOVA is robust against deviations from normality (Glass, Peckham, & Sanders, 1972; Harwell, Rubinstein, Hayes, & Olds, 1992; Schmider, Ziegler, Danay, Beyer, & Bühner, 2010). Where appropriate, ANOVAs were followed up with post-hoc t-tests. All reported correlations used Pearson’s correlation coefficient. Correlations were compared using Fisher’s *Z* transformation. Exploratory correlations between sleep variables and intrusion measures were also conducted.

## Results

### Sleep deprivation impairs the suppression of intrusive thoughts

First, we analysed how well participants controlled intrusive thoughts and investigated whether this was affected by sleep deprivation. Overall intrusion scores were robustly higher for ‘Think’ trials (M = 92.76% of trials, SEM = 1.38%) compared to ‘No-Think’ trials (M = 28.87% of trials, SEM = 2.52%) [*t*(58) = 19.73, *p* < .001, *d_z_* = 2.57], confirming that participants largely succeeded in suppressing thoughts about the scenes associated with red-framed faces. Because our primary goal was to examine intrusions for scenes that participants attempted to suppress, our analyses focus on ‘No-Think’ trials hereinafter, unless otherwise stated. Note that intrusion proportion scores for ‘No-Think’ trials on the mock TNT task did not differ between groups [sleep group: M = 41.38%, SEM = 3.76%; sleep deprivation group: M = 42.22%, SEM = 2.62%; *t*(57) = 0.19, *p* = .854], suggesting that any between-group differences in intrusion proportion for the TNT assessment phase proper were caused by our sleep/wake manipulation.

Of key interest is whether depriving participants of sleep disrupted their ability to stop unwanted thoughts from coming to mind, given reminders. Critically, sleep-deprived participants reported more intrusions than well-rested individuals [*F*(1,57) = 5.55, *p* = .022, *η_p_*^2^ = .09], demonstrating that sleep deprivation impairs memory control (Figure 2a). Strikingly, the sleep deprivation group suffered a proportional increase in intrusions of nearly 50% relative to the sleep group, revealing how deficient control may be a pathway to hyper-accessible thoughts. Participants overall showed decreasing intrusions across TNT trial blocks [*F*(3.16,180.33) = 29.81, *p* < .001, *η_p_*^2^ = .34; Greenhouse-Geisser corrected] (Figure 2b), indicating that repeatedly inhibiting retrieval was, in general, highly effective at stopping memories from intruding on future trials. Intrusion rates were comparable for negative and neutral scenes [*F*(1,57) = 0.11, *p* = .738], and the detrimental impact of sleep deprivation on intrusion rate did not vary with the valence of scenes [all *p* > .05].

**Fig. 2.**
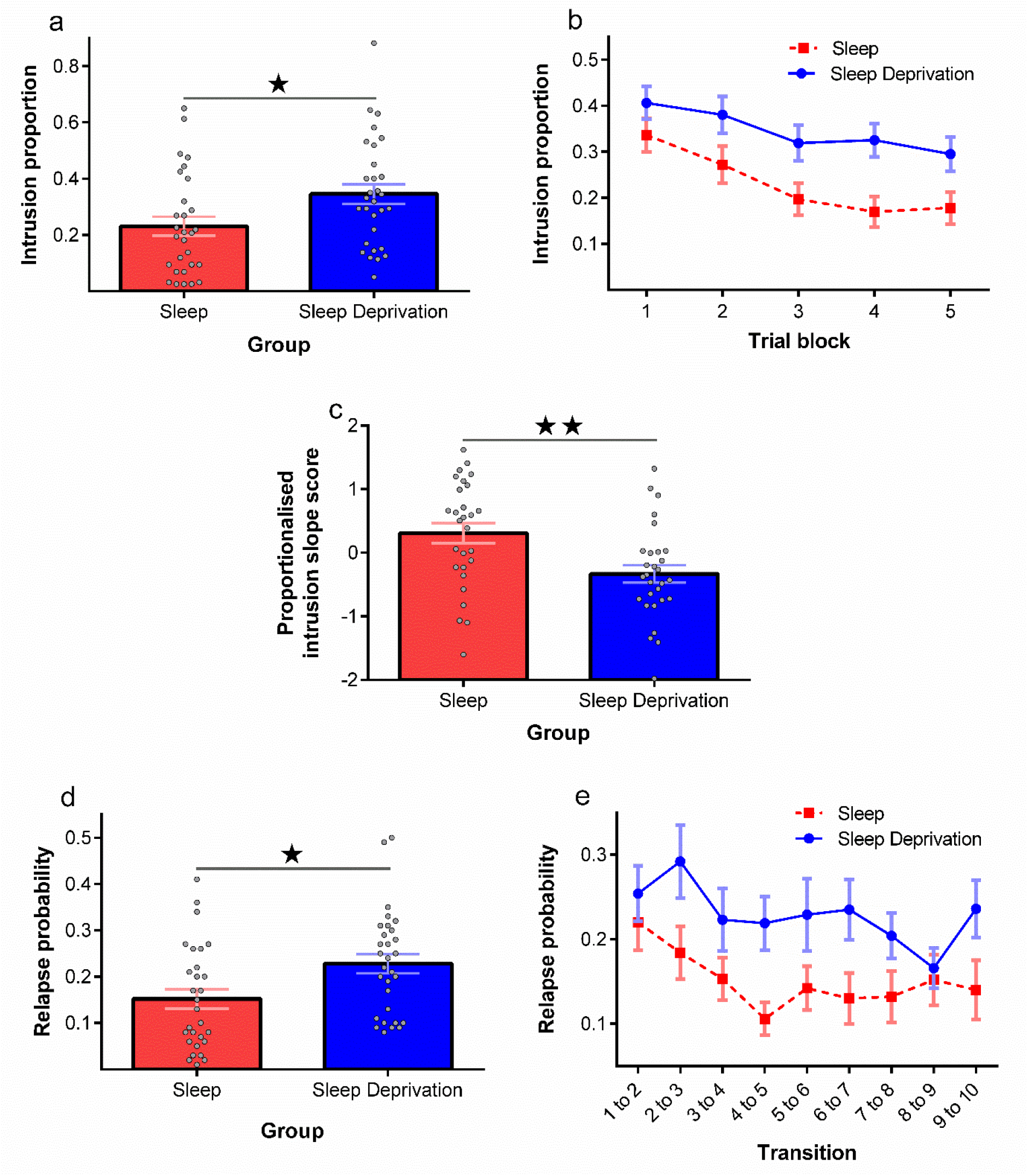
Sleep deprivation impairs the suppression of intrusive thoughts. (a) Intrusion proportion (i.e. proportion of ‘No-Think’ trials for which participants reported awareness of the associated scene), averaged across TNT blocks. (b) Intrusion proportion in each TNT block. (c) Proportionalised intrusion slope scores (i.e. the rate at which intrusions declined across the five TNT blocks for ‘No-Think’ items, divided by the number of intrusions in the first TNT block). Greater slope scores reflect more effective downregulation of intrusions across trial blocks. (d) Relapse probability (i.e. proportion of ‘No-Think’ trials where successful control on repetition *N* was followed by failed control on repetition *N*+1), averaged across trial transitions. (e) Relapse probability for each trial transition. Grey dots represent individual participants; data are shown as mean ± SEM; ★ represents *p* ≤ .05; ★★ represents *p* ≤ .01.

Although suppression robustly reduced intrusions over blocks across all participants, we were also interested in examining whether sleep deprivation altered the rate at which suppression attempts led intrusions to decline. Consistent with deficient inhibitory control, depriving participants of sleep significantly disrupted their ability to downregulate intrusive thoughts over blocks compared to participants who had slept, as reflected in lower intrusion slope scores [*F*(1,55) = 9.38, *p* = .003, *η_p_*^2^ = .15; Figure 2c]. Slope scores did not differ between negative and neutral scenes [*F*(1,55) = 0.08, *p* = .777], and the intrusion slope deficit in the sleep deprivation group did not vary with scene valence [*p* > .05].

We next considered whether sleep-deprived participants were more vulnerable to intrusion relapses for scenes that they had previously suppressed, as compared to participants that had slept. Sleep-deprived participants exhibited significantly higher relapse probability scores than did the sleep group [*F*(1,57) = 7.08, *p* = .010, *η_p_*^2^ = .11; Figure 2d], indicating that after initially gaining control over an intrusive thought, they suffered intrusion relapses more than participants who had slept. As with intrusions overall, relapses also became less frequent across trial transitions [*F*(6.39,364.12) = 2.49, *p* = .020, *η_p_*^2^ = .04; Greenhouse-Geisser corrected; Figure 2e]. Relapse probability was comparable for negative and neutral memories [*F*(1,57) = 0.07, *p* = .800], and sleep deprivation affected negative and neutral relapses to a similar degree [*p* > .05].

### Sleep deprivation nullifies affect suppression

Although the foregoing findings show that sleep deprivation disrupts intrusion control, they do not address whether losing sleep alters how suppression affects emotional responses. Recent work indicates that successfully suppressing negative scenes renders them less aversive when they are later re-encountered (Gagnepain et al., 2017). Building on this finding, we tested whether sleep deprivation affected the relationship between retrieval suppression and our behavioural index of affect suppression. Behavioural affect suppression refers to the difference in affect ratings of scenes between the first (pre-delay and TNT) and second (post-delay and TNT) session. Positive affect suppression scores indicate that participants felt less negative about the scenes in the second affect evaluation task.

Consistent with earlier reports using this procedure (Gagnepain et al., 2017), we found that, across all participants, successful retrieval suppression (i.e. fewer intrusions) was associated with greater affect suppression for negative scenes [*r* = - .32, *p* = .015; Figure 3a]. In other words, greater success at suppressing retrieval of aversive memories predicted a larger change in their perceived affect, rendering them less negative when participants next encountered the scenes. We observed no such relationship for neutral ‘No-Think’ scenes, or ‘Think’ scenes of either valence type [all *p* > .05]. The relationship between intrusion control and affect suppression did not differ significantly between the sleep [*r* = -.33, *p* = .079] and sleep deprivation groups [*r* = -.17, *p* = .367; *Z* = 0.62, *p* = .533].

**Fig. 3.**
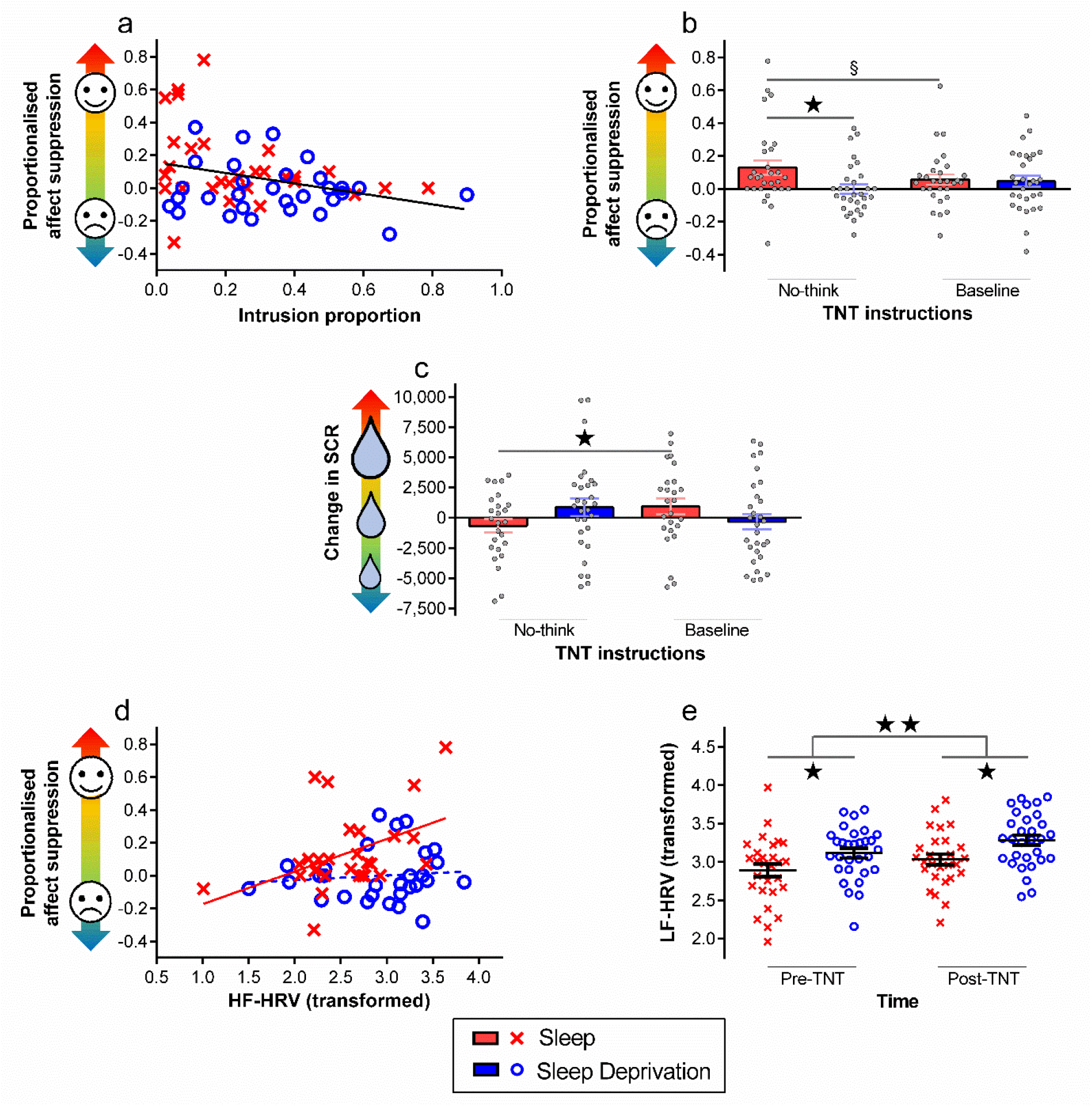
Sleep deprivation influences affect suppression and heart rate variability. (a) Correlation between intrusion proportion and affect suppression scores for negative ‘NoThink’ scenes. (b) Affect suppression scores for negative scenes. (c) Change in skin conductance response (dSCR) towards negative scenes. (d) Correlations between log-transformed resting HF-HRV (pre-TNT) and affect suppression for negative ‘No-Think’ scenes. (e) Mean log-transformed LF-HRV in each group. Stars represent main effects of group (sleep, sleep deprivation) and time (pre-TNT, post-TNT). Data points represent individual participants; data are shown as mean ± SEM; § represents *p* ≤ .10; ★ represents *p* ≤ .05; ★★ represents *p* ≤ .001.

Having found evidence of a general relationship between intrusion control and affect suppression specifically for negative ‘No-Think’ scenes, we next tested whether sleep deprivation undermines suppression-induced regulation of negative affect for aversive scenes. We observed a significant interaction between TNT Condition (‘Baseline’, ‘No-Think’) and Group [*F*(1,57) = 5.38, *p* = .024, *η_p_*^2^ = .09]. Affect change scores for aversive ‘No-Think’ scenes were greater in the sleep group than the sleep deprivation group [*t*(57) = 2.56, *p* = .013, *d* = 0.67; Figure 3b] supporting the view that the affective benefits of negative thought suppression were impaired after sleep loss. Note that we observed no such difference for negative ‘Baseline’ scenes [*t*(57) = 0.24, *p* = .809, *d* = 0.06], suggesting that the effect for negative ‘No-Think’ scenes was not driven by between-group differences in mood. Interestingly, in the sleep group, affect change scores for negative ‘No-Think’ scenes were more positive than those for negative ‘Baseline’ scenes, although this difference did not reach statistical significance [*t*(28) = 1.76, *p* = .089, *d_z_* = 0.33]. We observed no such difference in the sleep deprivation group [*t*(29) = 1.50, *p* = .145, *d_z_* = 0.27].

The foregoing findings indicate that suppression reduced the perceived valence of suppressed negative scenes. However, they do not speak to whether suppression alters sympathetic arousal, and whether sleep loss impacts this form of regulation. To probe this question, we analysed dSCR scores, which refer to the difference in SCRs across sessions arising for negative ‘Baseline’ and ‘No-Think’ scenes. Negative dSCR scores reflect a decrease in SCRs towards the scenes in the second affect evaluation task.

As with our behavioural index of affect suppression, we observed a significant interaction between TNT Condition (‘Baseline’, ‘No-Think’) and Group on dSCR scores [*F*(1,52) = 4.05, *p* = .049, *η_p_*^2^ = .07; Figure 3c]. Participants who slept exhibited a decrease in dSCR towards the negative ‘No-Think’ scenes compared to ‘Baseline’ scenes [*t*(24) = 2.07, *p* = .050, *d* = 0.41]. This finding is consistent with our behavioural affect suppression data, and further supports the view that successfully suppressing unpleasant memories alleviates their associated emotional charge. Sleep-deprived participants, who were less competent at controlling intrusions, exhibited comparable dSCR scores for ‘Baseline’ and ‘No-Think’ scenes [*t*(28) = 1.08, *p* = .290]. These findings suggest that the affective benefits of retrieval suppression for rested participants were not confined to subjective reports of perceived affect, but also arose for indices of psychophysiological emotional arousal. Sleep deprivation abolished the affective benefits of suppression, and this was evident in both our behavioural and psychophysiological measures. Interestingly, we observed no reliable correlation between individual differences in our behavioural and psychophysiological affect suppression measures for negative ‘No-Think’ scenes, suggesting that these indices tap partially distinct processes [*r* = 0.09, *p* = .522]. The observed change in psychophysiological reactivity for ‘No-Think’ scenes in the sleep group likely reflects diminished sympathetic nervous system responses to suppressed stimuli, possibly initiated by the impact of retrieval suppression on activity in the amygdala (Gagnepain et al., 2017).

### Heart rate variability is linked to affect suppression and fatigue

Inhibitory control over cognition is linked to HF-HRV (Gillie et al., 2014). This finding led us to investigate whether the degree of affect suppression that a given participant could achieve is linked to their HF-HRV, and whether any relationship is altered by sleep loss. Indeed, higher HF-HRV (collected prior to the TNT phase) predicted greater affect suppression scores for negative ‘No-Think’ scenes in the sleep group [*r* = .44, *p* = .021; Figure 3d]. This relationship was absent in sleep-deprived participants, suggesting that sleep deprivation may reduce the involvement of control processes related to HF-HRV [*r* = .11, *p* = .577; *Z* = 1.30, *p* = .097]. We observed the same pattern using the HF-HRV measure collected after the TNT phase, though to a lesser degree (sleep group [*r* = .34, *p* = .077]; sleep deprivation group [*r* = .07, *p* = .724; *Z* = 1.03, *p* = .152]). In contrast, we found no relationship between HF-HRV (pre-TNT or post-TNT) and affect changes for negative ‘Think’ or ‘Baseline’ scenes, or for neutral scenes in any TNT condition, for either group [all *p* > .05]. We found no relationship between HF-HRV and overall intrusion proportion for ‘No-Think’ scenes [both groups *p* > .05]. The LF-HRV component was not associated with affect suppression or overall intrusion proportion in any TNT condition for either group [all *p* > .05].

LF-HRV has been related to both sleep deprivation (Zhong, 2005) and mental fatigue (Egelund, 1982; Tran et al., 2009; Zhao, Zhao, Liu, & Zheng, 2012). Confirming this relationship, the sleep deprivation group exhibited higher LF-HRV than the sleep group [*F*(1,56) = 6.71, *p* = .012, *η_p_*^2^ = .11; Figure 3e]. Along similar lines, we found significantly higher LF-HRV following the TNT phase than we did preceding it [*F*(1,56) = 17.96, *p* < .001, *η_p_*^2^ = .24], reflecting task-induced fatigue, given the ~40 min duration of the TNT task (van Schie & Anderson, 2017). We observed no interaction between factors [*p* > .05].

Interestingly, performing the TNT task also affected HF-HRV: HF-HRV was higher after the TNT task than before it [*F*(1,56) = 19.62, *p* < .001, *η_p_*^2^ = .26]. An interaction between Time and Group [*F*(1,56) = 9.39, *p* = .003, *η_p_*^2^ = .14] drove this difference. Whereas completing the TNT task elevated HF-HRV in the sleep group [*t*(27) = 5.54, *p* < .001, *d_z_* = 1.05], it did not in the sleep deprivation group [*t*(29) = 0.93, *p* = .358, *d_z_* = 0.17], primarily because HF-HRV was already high at the outset in sleep-deprived participants. Overall, the groups did not differ reliably in HF-HRV [*F*(1,56) = 2.26, *p* = .138]. These data suggest that fatigue may increase HF-HRV, but less reliably than LF-HRV.

## Discussion

Our findings indicate that sleep deprivation substantially increases people’s vulnerability to unwanted memories intruding into conscious awareness when they confront reminders. Despite exhibiting comparable intrusion control ability before the overnight interval, sleep-deprived participants exhibited a near fifty percent proportional increase in intrusions relative to participants that had slept. Moreover, sleep deprivation diminished the cumulative benefits of retrieval suppression for downregulating subsequent intrusions. Even after sleep-deprived participants initially gained control over unwanted memories and prevented them from intruding, they were consistently more susceptible to relapses when reminders were confronted again later, as compared to rested individuals.

Deficient memory suppression following sleep loss might arise from dysfunction of the neural networks that govern inhibitory control. Retrieval suppression engages the rDLPFC, which is thought to downregulate recollection-related activity in MTL areas via inhibitory, top-down mechanisms (Benoit et al., 2015; Gagnepain et al., 2017; Levy & Anderson, 2012). The functional integrity of the rDLPFC may be particularly vulnerable to sleep loss (Mazur et al., 2002). Functional connectivity from the medial PFC (mPFC) to MTL appears to be disrupted in sleep-deprived individuals when viewing negative emotional images, potentially compromising a pathway of inhibitory control over affect (Yoo et al., 2007). Moreover, prolonged sleep restriction impairs performance on attention and working memory tasks that rely on prefrontal engagement (Chee & Choo, 2004; Frenda & Fenn, 2016; Lim & Dinges, 2008; Thomas et al., 2000). In the current study, sleep deprivation may have disrupted functional interactions between the rDLPFC (and possibly mPFC) and MTL structures such as the hippocampus and amygdala during retrieval suppression, impairing inhibitory control over memory and affect; increasing intrusions, and decreasing affect suppression. Future research can employ functional brain imaging to address these hypotheses.

Importantly, sleep deprivation also disrupted people’s ability to reduce the affective content of intruding thoughts through retrieval suppression. Consistent with earlier research (Gagnepain et al., 2017), we first confirmed that the more successful people are at controlling unpleasant intrusions, the greater their behavioural affect suppression for the suppressed scenes. Extending this finding, we showed that, in rested participants, affect suppression effects arose in psychophysiological reactivity, with suppressed scenes eliciting significantly reduced SCRs compared to scenes that were not suppressed, consistent with a reduction in sympathetic arousal. These behavioural and psychophysiological effects complement prior work showing that indices of affect suppression are predicted by downregulation of amygdala activity by prefrontal cortex during memory intrusions (Gagnepain et al., 2017), a process that could underlie both effects. Critically, sleep-deprived participants showed markedly less behavioural and psychophysiological affect suppression for negative ‘No-Think’ scenes, as compared to rested participants. These findings suggest that difficulties in engaging inhibitory control to regulate unpleasant intrusions after sleep loss lead to diminished affect regulation for the suppressed content.

The possibility that sleep deprivation compromised affect suppression by altering prefrontal control involvement receives indirect support from psychophysiological indices. Among participants who slept normally, higher HF-HRV predicted better affect suppression for negative ‘No-Think’ scenes; this relationship was not found, however, after sleep deprivation. Prior evidence links higher HF-HRV to better executive functioning (for review, see: Thayer & Lane, 2009), including memory control (Gillie et al., 2014). HF-HRV has been hypothesized to reflect the engagement of brain regions supporting cognitive and affective downregulation during memory inhibition, and in healthy participants blood flow in the right prefrontal cortex correlates with HF-HRV (Lane et al., 2009). Notably, however, in the current data, higher HF-HRV did not predict fewer intrusions for ‘No-Think’ scenes. The selective relationship of HF-HRV and affect suppression may indicate that this measure better indexes the efficiency of those components of the inhibitory control pathway that are uniquely tied to affect regulation (e.g. connectivity between mPFC and amygdala: Yoo et al., 2007). In the sleep deprivation group, extreme fatigue may have compromised these components, eliminating the relationship between HF-HRV and the emotional aftereffects of suppression, a possibility supported by research on the relationship between sleep deprivation, HF-HRV and cognitive performance (Quintana et al., 2017).

Prior research suggests that in healthy participants, acute sleep deprivation increases LF-HRV (Zhong, 2005). Replicating this work, our sleep-deprived participants exhibited higher LF-HRV compared to participants who slept. LF-HRV is produced by both sympathetic and parasympathetic systems, whereas HF-HRV reflects parasympathetic activity alone (Shaffer & Ginsberg, 2017). Given that HF-HRV was not altered by our sleep/wake manipulation, our results suggest that sleep deprivation elevates sympathetic arousal. Consistent with earlier work showing that mental fatigue can increase LF-HRV (Egelund, 1982; Tran et al., 2009; Zhao et al., 2012), completing our cognitively-demanding TNT task also elevated LF-HRV.

Negative scenes were no more intrusive than neutral scenes, despite a highly robust difference in perceived valence reported by our participants (see Methods). This finding might at first seem counterintuitive. However, following previous work which observed the same result (Gagnepain et al., 2017), we matched the initial training level of negative and neutral materials, which may have nullified the benefits of emotional arousal on encoding and consolidation (Canli, Zhao, Brewer, Gabrieli, & Cahill, 2000; Ritchey, Dolcos, & Cabeza, 2008). Failure to match initial training of negative and neutral materials may explain prior mixed findings regarding the relation between valence and forgetting; whereas some studies report enhanced suppression for negative relative to neutral stimuli (Depue, Banich, & Curran, 2006; Lambert, Good, & Kirk, 2010), others observed no difference (van Schie, Geraerts, & Anderson, 2013) or even the opposite effect (Nørby, Lange, & Larsen, 2010).

Although impaired memory control following sleep deprivation is a valid interpretation of our data, another possibility relates to overnight memory processing. Previous work has shown that sleep promotes the consolidation of procedural skills (for review, see: Stickgold, 2005). Accordingly, intrusion control may have been impaired in the sleep deprivation group as these participants did not have an opportunity for sleep-associated consolidation of the mnemonic suppression skills learned during the evening TNT practice. Although this interpretation is possible, participants were only given very brief practice with the TNT task (12 ‘No-Think’ trials on 6 filler items) in the first session, making it an unlikely source of the substantial sleep advantage in intrusion control that we observed. Nevertheless, future work should seek to distinguish the relative contributions of procedural memory consolidation and disrupted inhibitory control to greater intrusiveness after sleep deprivation.

Prior work suggests that sleep curtails spontaneous intrusive thoughts following trauma. Indeed, sleeping soon after exposure to a traumatic film clip reduces the number of spontaneous intrusions relating to that clip in the following week (Kleim, Wysokowsky, Schmid, Seifritz, & Rasch, 2016). Furthermore, greater sleep disturbance following a road traffic accident predicts more frequent spontaneous, accident-related thoughts during the first week post-trauma (Luik, Iyadurai, Gebhardt, & Holmes, 2019). Although these studies associate sleep loss with variations in intrusion frequency following adverse events, they do not address the mechanistic basis of these associations. Sleep following a traumatic film, for example, could reduce intrusions over the following week by enabling better memory control, or, alternatively, by facilitating sleep-related changes to the memory trace that make it less likely to intrude. As such, prior work has not established causal evidence concerning whether sleep deprivation impairs retrieval suppression of specific, unwanted thoughts. Because we used a procedure that demands the suppression of aversive memories in response to reminders, the current study pointedly targets the causal status of sleep loss in compromising inhibitory control over thought.

Our findings may help to explain the relationship between sleep disturbance and vulnerability to psychiatric conditions associated with intrusive thoughts. Up to ninety percent of patients with PTSD (Maher et al., 2006) and MDD (Riemann et al., 2001) report recurrent sleep disturbances, which are major risk factors for disorder onset (Maher et al., 2006; Mendlewicz, 2009). Exactly *how* disturbed sleep contributes to these disorders remains elusive, however. Our data raises the possibility that poor intrusion control may help bridge the gap between disturbed sleep and psychiatric symptoms: insufficient sleep might increase memory intrusions, whilst also nullifying the benefits of retrieval suppression for regulating affect. The onset of intrusive thoughts and affective dysfunction following bouts of poor sleep could create a vicious cycle, whereby upsetting intrusions and emotional distress exacerbate sleep problems (Talamini, Bringmann, de Boer, & Hofman, 2013), inhibiting the sleep needed to support recovery.

Besides PTSD and MDD, our findings might have implications for understanding other disorders linked to sleep disturbances, such as obsessivecompulsive disorder (OCD) and schizophrenia (Coles, Schubert, & Sharkey, 2012; Monti & Monti, 2005; Paterson, Reynolds, Ferguson, & Dawson, 2013). Acute sleep deprivation can induce dissociative symptoms in healthy individuals (Giesbrecht, Smeets, Leppink, Jelicic, & Merckelbach, 2007; Van Heugten-Van Der Kloet, Giesbrecht, & Merckelbach, 2015) and sleepiness exacerbates dissociative symptoms in depersonalization disorder (Simeon & Abugel, 2006). Observations such as these led researchers to hypothesise that sleep loss might fuel psychiatric disorders by promoting dissociation (van der Kloet, Merckelbach, Giesbrecht, & Lynn, 2012). Although dissociative symptoms are commonplace in OCD and schizophrenia, these conditions are also characterised by recurrent and persistent thought intrusions. Our data suggest that sleep loss could contribute to mental illness not only by fostering dissociative symptoms, but also by hindering unwanted thought suppression. It should be noted, however, that a night of total sleep deprivation, as employed in this study, is qualitatively different to the chronic sleep disturbances often observed in psychiatric conditions. Further studies are required to ascertain whether chronic sleep disturbance has similar consequences to acute sleep deprivation for intrusion control and affect suppression.

Evidence suggests that practice improves intrusion control in both real-life situations and via strategic intervention. For example, undergraduate students who report having experienced moderate childhood adversity exhibit greater memory control than those who report experiencing little to no adversity (Hulbert & Anderson, 2018). This finding suggests that prior experience in suppressing upsetting memories facilitates intrusion control even for new, unrelated memories, leading to greater resilience to adversities later in life. Moreover, providing MDD patients with a cognitive strategy to aid their intrusion control prior to performing a retrieval suppression task significantly facilitates memory inhibition (Joormann, LeMoult, Hertel, & Gotlib, 2009). These observations demonstrate that therapeutic strategies targeted at nurturing intrusion control could be utilized to prevent the development of psychiatric disorders in those who are at-risk on account of their sleep problems, such as insomnia sufferers (Baglioni et al., 2011).

We found that sleep deprivation markedly impairs the ability to prevent unwanted memories from entering conscious thought. Moreover, whereas suppressing negative memories after sleep renders them less subjectively and psychophysiologically aversive, poor intrusion control following sleep deprivation negates this affective benefit. Together with an increasingly specific understanding of the neural machinery of retrieval suppression (Anderson, Bunce, & Barbas, 2016; Anderson & Hanslmayr, 2014; Guo, Schmitz, Mur, Ferreira, & Anderson, 2017; Schmitz, Correia, Ferreira, Prescot, & Anderson, 2017), these findings point to an important neurocognitive mechanism linking sleep problems to the pathogenesis and maintenance of psychiatric conditions characterized by intrusive symptoms. Developing interventions that improve retrieval suppression in poor sleepers may be a promising avenue for averting the potentially pathogenic consequences of disordered control over distracting thoughts.

## Supporting information

Methodological Details S1

Methodological Details S2

Methodological Details S1

Recognition Task S1

## Author contributions

M.O.H, S.A.C., and M.C.A. designed the study; M.O.H and J.E.A. performed the experiments; M.O.H, S.S, M.C.A and S.A.C analysed the data; M.O.H wrote the manuscript; M.O.H, J.E.A, S.S, M.C.A and S.A.C revised the manuscript.

## Acknowledgments

This work was supported by a Medical Research Council Career Development Award (MR-P020208-1) to S.A.C, and a Medical Research Council grant (MC-A060-5PR00) to M.C.A. The authors are grateful to Rhiannon Pearce, Tomas Vaitkus and Hanna Weiers for their assistance with data collection, and to two anonymous reviewers for their helpful comments on an earlier version of this manuscript.

## Declaration of interests

The authors declare no competing interests.

## Data availability

Study data are available via the following link: https://osf.io/95dbh/

## Notes

#### Summary of Updates

Manuscript updated to address comments made by two anonymous reviewers. Manuscript made significantly shorter, in part by moving entire sections to Supplementary Material.

