## Supplementary material for "Losing Control: Sleep Deprivation Impairs the Suppression of Unwanted Thoughts": Methodological Details S1

Electrodermal activity was recorded using a BIOPAC MP36R data acquisition system and AcqKnowledge (ACQ) 4.4.1 software (sampling rate = 2KHz). During the affect evaluation tasks, E-Prime-triggered square pulse outputs were transmitted to the MP36R unit via a BIOPAC STP35A interface enabling precise alignment of each stimulus onset to the skin conductance response (SCR) data. Two BIOPAC EL507 disposable adhesive electrodes were attached to the fingertips of the index and middle fingers of the non-dominant hand. The data were imported and preprocessed using PsPM (version 4.0.2; Bach & Friston, 2013).

A unidirectional first-order Butterworth high-pass filter with cut-off frequency 0.05 Hz was used to filter the data to account for the change of baseline activity during the duration of the recording sessions. The time series averaged over corresponding trials for each experimental condition (e.g. ‘No-Think’, negative valence) were then extracted, for each subject, for each of the two sessions (pre-TNT and post-TNT; TNT = Think/No-Think). An average ‘session-specific’ skin conductance level (SCL) was computed for each subject for each session. SCL was the skin conductance value measured for the first second after the presentation of the stimuli, averaged across all conditions. This first 1 s period after stimulus presentation is widely considered to be the ‘latency’ period for event-related evoked SCRs (Bach, Flandin, Friston, & Dolan, 2010; Braithwaite, Watson, Robert, & Mickey, 2013; Lim et al., 1997). This ‘baseline’ SCL was then subtracted from the rest of the measured skin conductance activity, which was deemed to belong to a canonical evoked SCR, taken for the entirety of the time that the stimulus was presented on screen during a trial, after excluding the first second (i.e. 5.5 seconds). The area under the curve for each condition was then computed for each subject, for each of the two sessions. This is akin to an analysis approach shown previously for spontaneous skin conductance fluctuations, except here we have used it to analytically quantify event-related evoked SCRs (Bach, Friston, & Dolan, 2010). The average SCRs elicited by scenes in each TNT condition and valence category were used to calculate the difference in SCRs across sessions (dSCR; SCR post-TNT – SCR pre-TNT). The same analysis pipeline was used for both the sleep and the sleep deprivation groups. Data from 2 participants were unavailable due to technical issues (sleep group *n*=1; sleep deprivation group *n*=1). Furthermore, we excluded data from 3 participants in the sleep group who were SCR non-responders.
