## Supplementary material for "Losing Control: Sleep Deprivation Impairs the Suppression of Unwanted Thoughts": Methodological Details S2

Electrocardiography (ECG) was recorded using a BIOPAC MP36R data acquisition system and AcqKnowledge (ACQ) 4.4.1 software (sampling rate = 2KHz). Three BIOPAC EL503 ECG electrodes were attached to the midline of the left and right clavicle and the lower left rib. ECG was recorded for eight successive minutes. The first 2 min and last 1 min of each recording was discarded, and heart rate variability (HRV) was calculated for the remaining five successive minutes. The ECG signal was analysed offline using ACQ and Kubios Standard 3.0.2 software.

R-peaks were automatically detected using ACQ’s QRS detection algorithm and visually inspected for accuracy. Peaks that the algorithm missed were inserted manually. The interbeat-interval time series was then imported to Kubios for analysis. To obtain frequency-domain-specific indices of HRV, we used autoregressive estimates of low-frequency (0.04-0.15 ms^2^/Hz) and high-frequency (0.15-0.40 ms^2^/Hz) power. Autoregressive algorithms are generally preferable to Fourier transform based algorithms for spectral analysis of HRV (Thayer, Hansen, & Johnson, 2008), partly because they have better spectrum resolution when using short data frames (Miranda et al., 2012). In keeping with previous research (Gillie, Vasey, & Thayer, 2014; Park, Vasey, Van Bavel, & Thayer, 2014), values of low-frequency HRV (LF-HRV) and high-frequency HRV (HF-HRV) were transformed logarithmically (base 10). One participant in the sleep group exhibited atypical ECG patterns, and was thus removed from HRV analyses.
