## Supplementary material for "Losing Control: Sleep Deprivation Impairs the Suppression of Unwanted Thoughts": Methodological Details S1

Methodological Details S3

Sleep monitoring was carried out using an Embla N7000 polysomnography (PSG) system (Embla Systems, Broomfield, CO, USA). Gold-plated electrodes were attached using EC2 electrode cream after the scalp was cleaned with NuPrep exfoliating agent. Scalp electrodes were attached at eight standard locations according to the international 10-20 system (Homan, Herman, & Purdy, 1987): F3, F4, C3, C4, P3, P4, O1, and O2, each referenced to the contralateral mastoid (A1 or A2). Left and right electrooculogram, left, right and upper electromyogram, and a ground electrode (forehead) were also attached. All electrodes were verified to have a connection impedance of < 5 kΩ. All signals were digitally sampled at a rate of 200 Hz.

Sleep data was divided into 30 s epochs and scored as wakefulness, N1 sleep, N2 sleep, slow-wave sleep (SWS) or rapid eye movement (REM) sleep according to standardised criteria (Iber, Ancoli-Israel, Chesson, & Quan, 2007), using RemLogic 3.4. PSG data was unavailable for 4 participants due to reference electrodes becoming detached during the night.
