## Supplementary material for "Losing Control: Sleep Deprivation Impairs the Suppression of Unwanted Thoughts": Recognition Task S1

At the end of the experiment, memory for all of the face-scene pairs was tested using a recognition task. This task was included to ensure that participants had retained knowledge of the face-scene pairs across the overnight interval.

On each trial, participants viewed a single face, together with two scenes: one that was paired with the face and another which featured in the experiment but was not paired with this particular face. Participants were instructed to indicate which scene was paired with the face via key press within 5 s. We asked participants to make this response as quickly and accurately as possible. The trial terminated once a response had been provided or the time limit expired, before the next trial began.

Recognition accuracy was calculated as the proportion of face-scene associations that were correctly identified. Data were analysed using a 3 (TNT Condition: ‘Baseline’/’No-Think’/’Think’) x 2 (Valence: Negative/Neutral) x 2 (Group: Sleep/Sleep Deprivation) mixed ANOVA.

Recognition accuracy was very high in both groups [sleep group: M = 97.70%, SEM = 0.54%; sleep deprivation group: M = 96.32%, SEM = 0.71%], suggesting that knowledge of the face-scene pairs was well-retained across the overnight interval. There was no statistically significant difference in recognition performance between groups [*F*(1,57) = 2.37, *p* = .129], and there was no significant main effect of TNT Condition [*F*(1.80,102.54) = 2.69, *p* = .078, *Greenhouse-Geisser corrected*] or Valence [*F*(1,57) = 0.36, *p =* .551], and there were no significant interactions [all *p* > .05].
