## Supplementary material for "Losing Control: Sleep Deprivation Impairs the Suppression of Unwanted Thoughts": Table S1

**Table S1.** Sleep stage data

| N1 (min; %) | 8.02 ± 1.07 | 1.96 ± 0.26 |
| --- | --- | --- |
| N2 (min; %) | 218.14 ± 6.76 | 53.56 ± 1.25 |
| SWS (min; %) | 107.34 ± 3.85 | 26.47 ± 0.93 |
| REM (min; %) | 73.00 ± 3.87 | 18.00 ± 0.95 |
| TST (min) | 406.50 ± 6.78 | - |
| Sleep efficiency (%) | 97.26 ± 0.49 | - |

Data presented as mean ± SEM. Sleep efficiency refers to the proportion of time spent asleep between sleep onset and final awakening. Abbreviations: N1, N2, stages of non-REM sleep; SWS, slow-wave sleep; REM, rapid eye movement sleep; TST, total sleep time.
